# MettleRNASeq: Complex RNA-Seq Data Analysis and Gene Relationships Exploration Based on Machine Learning

**DOI:** 10.1101/2025.05.06.652387

**Authors:** Samella Salles, Otávio Brustolini, Luciane Ciapina, Kary Ocaña

## Abstract

Typical differential gene expression (DGE) analysis might struggle when RNA-Seq datasets possess characteristics that hinder the power of statistical analyses and the obtention of accurate conclusions, such as a limited number of replicates and high variability. We present MettleRNASeq, a robust alternative for complex RNA-Seq data analysis that integrates machine learning techniques - a tailored classification approach, association rule mining, and complementary correlation analysis - to accurately identify key genes that distinguish experimental conditions and emphasize gene relationships. This approach provides full control over critical parameters, making it versatile for transcriptomic analyses and enhancing the comprehension of disease mechanisms, treatments, and their progression. MettleRNASeq was applied for the analysis of complex radiotherapy datasets. While popular DGE tools showed an inability to accurately differentiate the distinct radiotherapy treatments, MettleRNASeq effectively and consistently indicated relevant genes for condition discrimination and identified meaningful gene relationships related to radiotherapy, highlighting condition-specific and shared gene relationships. MettleRNASeq is implemented as an R package and available on GitHub at https://github.com/SamellaSalles/MettleRNASeq.

## 1. Introduction

RNA-Seq has transformed transcriptomics by enabling comprehensive gene expression analysis. A typical RNA-Seq analysis identifies differentially expressed genes (DEG) through statistical methods, predominantly relying on p-values. (Chavan-Gautam et al., 2017; Skerrett-Byrne et al., 2023; Yoo et al., 2018; Lowe et al., 2017). However, this approach may struggle when datasets have characteristics that hinder statistical power. High variability and a limited number of replicates, among others, can all affect DGE’s result reliability, yielding erroneous conclusions (Lu and Belitskaya-Levy, 2015; Lakens, 2021; Kitchin and Lauriault, 2015; Xu et al., 2023; Ching et al., 2014; Todd et al., 2016; Akobeng, 2016; Yu et al., 2017; Su et al., 2020).

Integrating Machine Learning (ML) into RNA-Seq analysis provides a promising alternative. Classification methods help identify gene sets that distinguish experimental conditions (Arowolo et al., 2021; Chen and Dhahbi, 2021). However, the high-dimensional nature of RNA-Seq data, where genes (features) far outnumber samples (observations), poses challenges for condition discrimination. While studies suggest techniques like ensemble methods and feature selection to improve classification (Jabeen et al., 2017; Tan et al., 2014; Arowolo et al., 2021), standard approaches can still struggle, particularly when the number of replicates is limited and variability is high (Cheng et al., 2024; Bishop and Nasrabadi, 2006).

Another relevant application of RNA-Seq data analysis is studying gene relationships, offering deeper insights into biological mechanisms across diseases, treatments, and their progression (Ruan et al., 2010; Lemoine et al., 2021). Beyond aiding biological interpretations, this relationship-focused analysis can help to extract meaningful information from complex datasets, which can be particularly difficult, presenting intricate biological contexts. Machine learning-based Association Rule Mining (ARM) and complementary Correlation Analysis (CA) can be employed to explore gene relationships, as ARM identifies conditional gene relationships by detecting “if-then” patterns (Chen et al., 2015; Gakii and Rimiru, 2021) and CA reveals the directionality and linearity (or not) of gene pair relations (Qian et al., 2022). Despite its potential, ARM is still underexplored in RNA-Seq analysis, with very few studies employing ARM (Gakii and Rimiru, 2021; Chen et al., 2015). Also, the application of CA is often limited to sample similarity studies (Koch et al., 2018) and comparison of different methods (Fonseca et al., 2014) or genes to biological mechanisms (Du et al., 2024; Chen et al., 2024), among others. Although correlation can be performed on specific gene relationships (Liu et al., 2024), it is often not focused on the comprehensive understanding of gene relationships related to specific conditions. To the best of our knowledge, no current works have exploited these three methods for RNA-Seq data analysis and for a thorough study of the biological mechanisms related to experimental conditions and their gene relations.

This work presents MettleRNASeq, a robust alternative for complex RNA-Seq analysis that integrates tailored classification with ARM and CA to discriminate experimental conditions and uncover gene relationships. MettleRNASeq enhances classification power by improving data representation and leveraging resampling, over-fitting, and testing. It also allows researchers to control key parameters throughout the analysis, making it adaptable to various transcriptomic studies. By integrating different machine learning techniques, MettleRNASeq provides a comprehensive understanding of biological processes, identifying key genes, pathways, and relationships that traditional methods might miss.

MettleRNASeq was evaluated on complex radiotherapy treatment datasets and compared to the traditional DGE analysis. While popular DGE tools showed an inability to accurately differentiate the distinct treatments, MettleRNASeq consistently identified relevant key genes for condition discrimination and indicated meaningful radiotherapy-related gene relationships. Furthermore, it captured condition-specific and shared gene relationships, emphasizing gene relations differentially regulated depending on the experimental condition.

## 2. Methodology

### 2.1 MettleRNASeq Approach

As illustrated in Figure 1, MettleRNASeq consists of data preparation steps followed by three key analyses enabling downstream biological studies: a tailored classification approach and relationship-focused methods, including correlation analysis and association rule mining.

**Figure 1.**
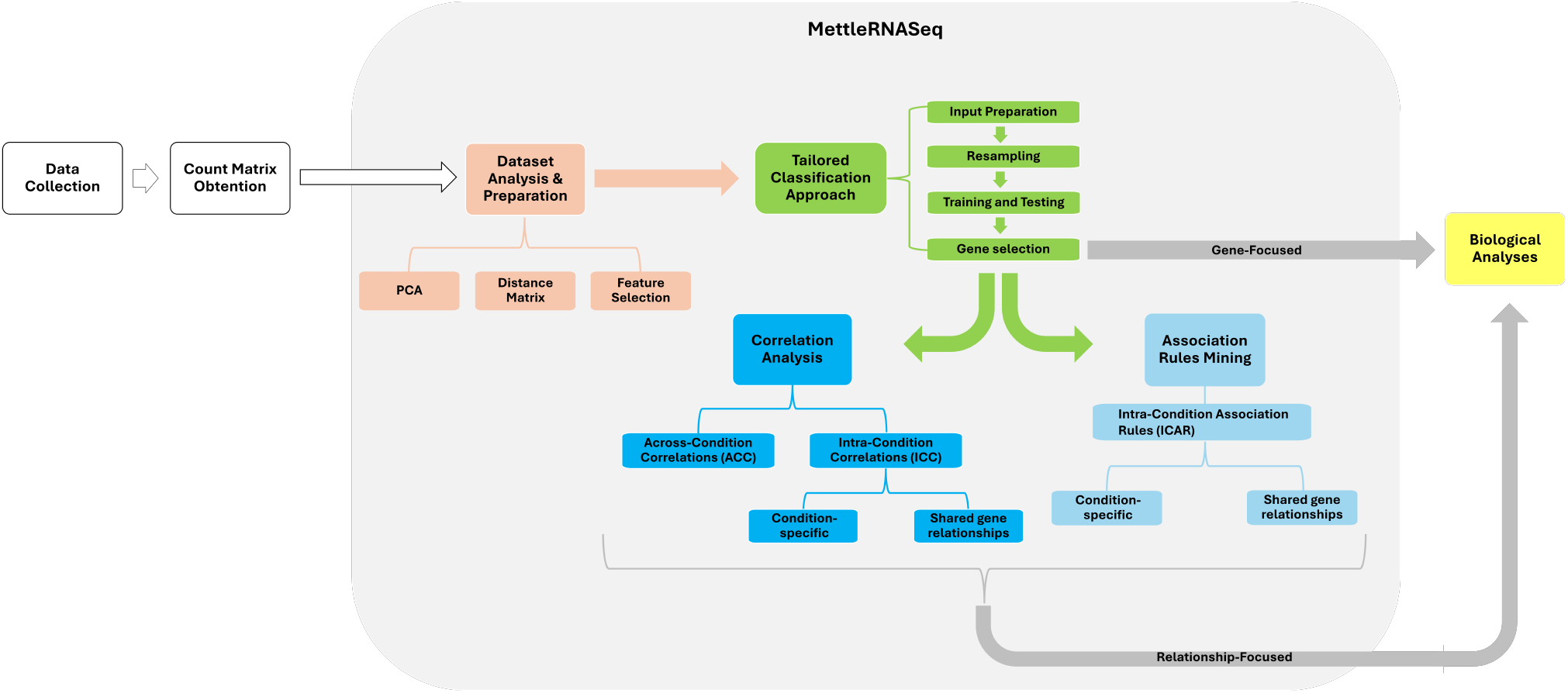
Overview of the MettleRNASeq approach. Data obtention steps are shown in the white rectangles. MettleRNASeq approach is illustrated in the light gray box, encompassing data preparation steps (in light pink) and the three key analyses: tailored classification (in green) and the gene relationship-focused correlation analysis and association rule mining (in blue tones). Biological analyses both gene and relationship-focused are shown in yellow.

#### 2.1.1 Dataset Analysis and Preparation

The dataset’s underlying structure is examined using Principal Component Analysis (PCA) and a distance matrix computation to assess data variability and sample similarities. Based on the PCA results, feature selection is performed to identify the genes that contribute most to the variance between the conditions of interest. Finally, the count matrix is filtered to retain only these relevant genes, forming the Refined Count Matrix (RCM) used for the following MettleRNASeq analyses.

#### 2.1.2 Tailored Classification Approach

This step identifies optimal genes for classifying the conditions under study, including Input Preparation, Resampling, Training and Testing, and Gene Selection, as detailed in Figure 2 and the following subsections.

**Figure 2.**
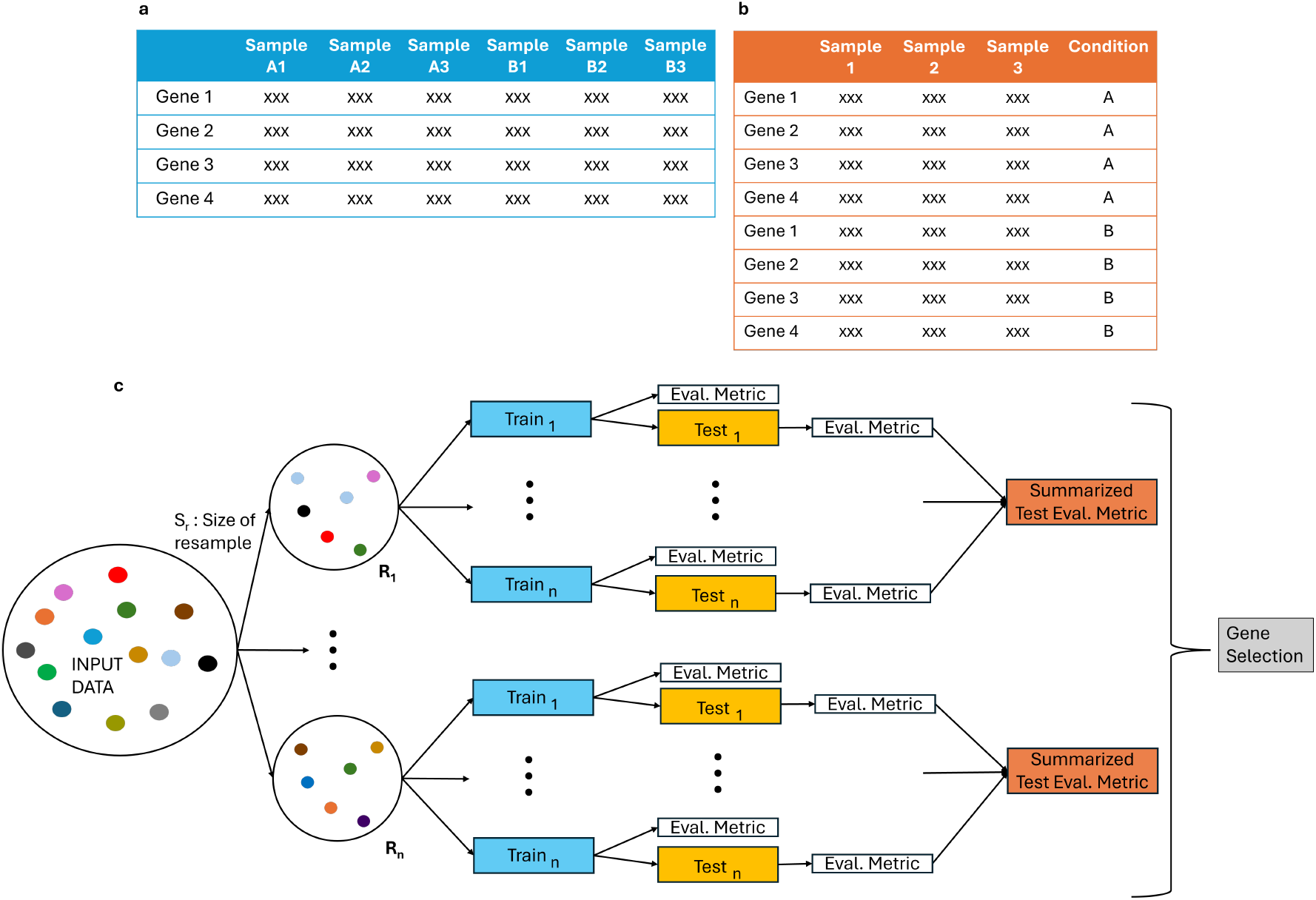
Overview of the tailored Classification Approach. (a) and (b) show the Input Preparation step, in which “xxx” represents the gene expressions. (a) illustrates a count matrix and (b) shows the input of the tailored classification approach after preparation, with “A” and “B” representing different conditions. (c) shows the steps after input preparation. The biggest circle represents the input data, where each colored dot illustrates a gene. The smallest circles (from *R*_1_ to *R*_*n*_) represent new resamples created during resampling, in which *S*_*r*_ symbolizes the resample size. Same colored dots (seen in *R*_1_) indicate that replacement in resampling is enabled. Training and Testing are displayed in the blue and yellow rectangles, respectively. The summarized test evaluation metrics are displayed in the orange rectangle. Finally, the gene selection step is shown in the gray rectangle.

##### 2.1.2.1 Input Preparation

MettleRNASeq restructures the RCM (Figure 2 a) so that gene expression values for each condition are stacked row-wise, ensuring that each gene is represented across all conditions. A ‘Condition’ column is appended to the matrix, labeling the conditions of interest for classification. This modified count matrix (Figure 2 b) serves as the input for the classification approach described in this work.

Since classification performs resampling across rows (where rows usually represent samples), this data restructuring enables the use of genes instead of samples. This strategy can be particularly advantageous for RNA-Seq data since the number of genes is much larger than samples, significantly increasing the number of resamples that can be generated during the resampling step, described in the following subsection. Especially in small sample size datasets, by leveraging genes for resampling, our classification approach enhances the representation of the conditions and identification of key genes that effectively discriminate between conditions.

##### 2.1.2.2 Resampling

The resampling step allows the choice of different resample sizes (*S*_*r*_), determining the number of genes randomly selected from the input data for all conditions, with or without replacement. When replacement is enabled, the same gene can appear multiple times in a resample, as illustrated by the light blue dots in *R*_1_ (Figure 2 c). The option to choose the *S*_*r*_ helps to determine the optimal number of genes for classifying conditions and to assess how much they differ. It also allows researchers to focus on a defined number of genes that best distinguish the conditions.

The number of resamples (*n*) is also configurable, allowing the exploration of distinct gene combinations. While a higher *n* is generally desirable, enabling the exploration of a wider range of gene combinations, a smaller *n* can still effectively represent the data and overcome resource limitations. For each combination of *S*_*r*_ and *n*, a table is generated listing the genes comprised in each resample (*R*_1_ to *R*_*n*_) of that resample size (*S*_*r*_).

##### 2.1.2.3 Training and Testing

Data partition generates multiple training and testing sample subsets, allowing the assessment of model performance across different data splits. A table is created to track which samples are assigned to each subset.

During training, the classification algorithms and evaluation metrics are chosen. Since some algorithms perform an internal test during training, this data split on the genes must be prevented by using the same data for both internal training and testing partitions. This allows classification on the defined number of genes and induces overfitting, making the model tailored to the given data (Bradshaw et al., 2023).

Testing is conducted on the distinct test subsets to evaluate model performance on unseen data. Test evaluation metrics (TEM) and a summarized TEM (STEM) for each resample are obtained, as illustrated in Figure 2. Finally, an evaluation table is produced for each combination of *S*_*r*_ and *n*, detailing the *R*_*n*_, training and testing subsets (*Train*_*n*_ and *Test*_*n*_), as well as their respective evaluation metrics and STEM. The STEMs for each of the combinations (*S*_*r*_ and *n*) can be compared to identify any trends in model performance.

##### 2.1.2.4 Gene selection

Resamples with evaluation metrics (TEMs or STEMs) above a defined threshold can be selected, and their genes aggregated and filtered based on uniqueness or recurrence to identify the best genes for condition discrimination. The RCM (Figure 2 a) can then be filtered to retain only these genes and used for downstream analysis.

#### 2.1.3 Gene Relationships Exploration

Correlation Analysis and Association Rule Mining are performed to explore the relationships of the genes selected (Figure 1). For that, the filtered RCMs are divided by condition into separate tables, referred to as the Condition Matrix (CM), allowing for condition-focused studies.

##### 2.1.3.1 Correlation Analysis

The CA is performed on the CMs, focusing on Across-Condition Correlations (ACC) and Intra-Condition Correlations (ICC). Methods such as Spearman, Kendall, or Pearson are chosen to allow capturing linear (Pearson) and non-linear (Spearman, Kendall) associations (Essam et al., 2022).

For ACC analysis, correlations are calculated for each gene across different conditions, revealing how gene expression patterns vary between experimental conditions. For ICC analysis, correlations are obtained within each condition for all gene pairs, uncovering condition-specific gene relationships. The comparison of ICC values allows the detection of shared gene pair relationships that remain consistent between conditions.

##### 2.1.3.2 Association Rule Mining

ARM is applied to the CMs to uncover conditional relationships (rules) between genes under specific conditions. First, the CM is converted into a Transaction class, in which each sample represents a transaction and each gene is categorized into three expression ranges (small, medium, and high), resulting in a total of *number of genes* × 3 possible items *per* transaction.

ARM algorithms such as Apriori or FP-Growth are employed (Lin et al., 2002). Key parameters, including the minimum and maximum rule length (i.e., the number of genes in a rule), support, and confidence thresholds, are carefully selected to balance the trade-off between capturing meaningful relationships and ensuring computational efficiency. The support threshold determines the minimum frequency at which an item (gene expression range) must appear across all transactions (samples) to be considered significant. The confidence threshold ensures that the identified rules have a high conditional probability of occurring, thereby enhancing the reliability of the findings (Lin et al., 2002).

Given the large number of rules typically generated by ARM, rule filtering is conducted to improve interpretability and focus on biologically relevant associations. The lift measure serves as a threshold, evaluating the strength of associations between genes and indicating meaningful conditional relationships when lift is greater than 1. Redundant rules - characterized by higher complexity, involving more genes but offering equal or lower predictive power than less complex rules - are removed during this filtering process. Another way to filter ARM results is by focusing on rules involving specific genes of interest, enabling targeted investigations (Lin et al., 2002).

Intra-Condition Association Rules (ICAR) are generated, capturing conditional relationships between genes and their respective ranges. Genes appearing in the same rules are clustered, forming sets that reveal coordinated associations under specific conditions. These clusters offer valuable insights into gene interactions and serve as a foundation for further biological analyses. By comparing ICARs, both condition-specific and shared gene relationships can be identified.

## 3. Experimental Evaluation

### 3.1 Case Study: Radiotherapy

We evaluated our approach using datasets from two different tissue types in radiotherapy studies of two modalities: Standard and the novel FLASH radiotherapy. While Standard damages cancer and healthy cells, FLASH delivers radiation at an ultra-high dose rate (*≥* 40 Gy/s), 400 times faster than Standard, achieving antitumor effectiveness comparable to that of Standard while reducing damage to healthy tissue - a phenomenon known as the FLASH effect (Chow and Ruda, 2024).

These two tissue datasets are an ideal case study for evaluating MettleRNASeq’s robustness and consistency across different biological environments for various reasons: The complexity of radiotherapy and cancer is well known, involving intricate mechanisms such as immune response, oxidative stress, DNA damage repair, cell proliferation, survival, and cell death (Swanton et al., 2024; Amundson, 2022; Chen and Kuo, 2017; Lu et al., 2022). Combined with tumor and radiation heterogeneity and tissue-specific radiation sensitivity, these factors complicate data analysis (Viana et al., 2011; Zhu et al., 2021; Macian, 2006; Cassidy et al., 2015). Moreover, distinguishing between FLASH and Standard radiotherapies becomes particularly challenging since both treatments share a radiation-based nature despite their differences in effects and dose rates (Lv et al., 2022; Lin et al., 2021; Matuszak et al., 2022; Velalopoulou et al., 2021). Finally, both datasets present a limited number of biological replicates *per* experimental condition.

#### 3.1.1 Data collection

We evaluated our approach using publicly available RNA-Seq datasets retrieved from the Gene Expression Omnibus (GEO) database at NCBI (Accession GSE173944). These data, generated by Velalopoulou et al. (2021), were obtained from a female mouse sarcoma model (*Mus musculus*) to investigate gene expression alterations in two tissues (skin and bone) in response to Proton Radiotherapy (PRT). Velalopoulou et al. (2021) examined two PRT modalities: Standard Proton Radiotherapy (S-PRT) and FLASH Proton Radiotherapy (F-PRT). Tissues were collected five days after irradiation to the right hind leg of the mouse. The three experimental conditions included: S-PRT-treated samples (Standard (S)); F-PRT-treated samples (FLASH (F)); Untreated controls (U). Each tissue dataset consisted of twelve samples comprising four biological replicates *per* condition.

#### 3.1.2 Count Matrix Obtention

FastQC (version 0.12.1) (Andrews et al., 2010) and MultiQC (version 1.17)(Ewels et al., 2016) were used for pre- and post-cleaning quality assessments. Data cleaning was performed with Cutadapt (version 4.5) (Martin, 2011) to remove low-quality reads (phred quality score < Q30), trim Illumina Universal Adapter sequences, and filter out short reads after trimming (<50 bp). STAR (version 2.7.11) (Dobin et al., 2013) was used to map the reads to the *Mus musculus* reference genome (GRCm39, Release M35) obtained from GENCODE (Frankish et al., 2023). The reads were counted with HTSeq-Counts Mode Union (version 2.0.4) (Anders et al., 2015) to obtain the count matrices for each combination of tissue (Bone and Skin), condition group (FLASH, Standard, and Untreated), and replicate (1 to 4).

R (version 4.4.1) packages were extensively used throughout the rest of this experimental evaluation and MettleRNASeq implementation. After the generation of the count matrices with HTSeq-Counts, they were merged into two dataframes, one *per* tissue, referred to as Tissue Count Matrix (TCM). The TCMs served as the basis for the MettleRNASeq approach, as illustrated in Figure 1.

#### 3.1.3 MettleRNASeq Approach

##### 3.1.3.1 Data Analysis and Preparation

Low-expression genes were filtered out from the TCMs, retaining only gene expressions ≥ 10 in all four replicates of each condition. The data were normalized using the popular and well-established Trimmed Mean of M-Values (TMM) method (Robinson and Oshlack, 2010)

Principal Component Analysis (PCA) (Wold et al., 1987) was performed using the *prcomp* function from the *stats* package (version 4.4.1) (R Core Team, 2024) with default settings (including “center=TRUE” and “scale=FALSE”). A three-dimensional PCA plot was generated using the *plot_ly* function from the *plotly* package (version 4.10.4) (Sievert, 2020).

The Euclidean distance matrix was computed using the *dist* function from the *stats* package and visualized as a heatmap using the *pheatmap* function from the *pheatmap* package (version 1.0.12) (R Core Team, 2024; Kolde, 2019).

Based on the PCA and distance matrix, the Untreated condition was filtered out of the TCMs, as it formed a distinct cluster and was the main contributor to overall data variation. A second PCA was then performed to capture the variation between the FLASH and Standard treatment conditions. Feature selection was applied to refine the TCMs, and principal components 1 to 4 were selected, together explaining over 95% of the variance. For each component, the top 100 genes with the highest contribution (i.e., rotation values) were chosen. After removing duplicates, a final set of 235 genes for Bone tissue and 186 for Skin was obtained and used to construct the RCMs for MettleRNASeq.

##### 3.1.3.2 Tailored Classification Approach

The RCMs were modified as described in the Input Preparation section in 2.1.2.1, with the Condition column stating either “FLASH” or “STANDARD” and converted into a factor with two levels. Resampling was performed with resample sizes (*S*_*r*_) of 10, 20, 30, 40, and 50 genes randomly selected from both conditions, with replacement enabled. The number of resamples (*n*) chosen were 100, 500, 1000, 5000, and 10000. Since the dataset presented four biological replicates *per* condition, data partitioning was performed using a leave-one-out approach, with one replicate set aside for testing. This resulted in four training-testing subsets: *Train*_1_ to *Train*_4_, where replicates 4, 3, 2, and 1 were excluded, respectively; and *Test*_1_ to *Test*_4_ formed by the corresponding left-out replicates, as illustrated in Figure 2.

The *Caret* R package (version 7.0.1) (Kuhn and Max, 2008) was used for classification using the *train* and *predict* functions for training and testing, respectively. The Random Forest algorithm (Breiman, 2001), a widely used ensemble method, was selected for training. Model performance was evaluated using the Accuracy metric. The internal test performed during the Random Forest training step was disabled, as described in section 2.1.2.3. The median of the TEMs was calculated to obtain the STEMs for each of the resamples created (Figure 2 c). Genes from the resamples with the highest median test accuracies (STEMs) were aggregated, and duplicate entries were removed to generate a final set of unique genes identified as the most effective for classifying the two conditions. Next, the CMs were constructed for each tissue (section 2.1.2.4) and utilized for the following gene relationship exploration analyses.

##### 3.1.3.3 Correlation Analysis

The Spearman method was used to capture monotonic relationships, including non-linear ones (Essam et al., 2022), with correlations computed using the *cor* function from the *stats* package (R Core Team, 2024) with default parameters.

For ICC, correlation matrices were visualized using the *corrplot* function from the *corrplot* package (version 0.95) (Wei and Simko, 2024). We applied the Angular Order of Eigenvectors (AOE) method through the *order* parameter, optimizing correlation matrices’ arrangement to highlight patterns and provide deeper insights into gene relationships.

Condition-specific and shared gene relationships were identified by calculating the absolute correlation difference between conditions, emphasizing both unique and conserved gene relations throughout conditions within the tissue.

##### 3.1.3.4 Association Rule Mining

We utilized the *arules* package (version 1.7.8) (Hahsler et al., 2005) to perform ARM. The CMs were converted into the Transaction class using the *as* function. Each condition presented four transactions (samples) and a total of *number of unique genes selected ×* 3 possible items (gene expression ranges) *per* transaction.

The widely used Apriori algorithm (Hegland, 2007) was employed via the *apriori* function. The minimal rule length was set to 2 items, ensuring that each rule included at least two genes. The maximal rule length was set to 10, the maximum allowed by the function to prevent excessively long run times. The support threshold of 0.5 was chosen to highlight frequent items appearing in at least half of the transactions while maintaining computational efficiency. The confidence threshold was set to the default value of 0.8, effectively filtering out rules with low conditional probability. We applied the *is*.*redundant* function to remove redundant associations, and only rules with a lift greater than 1 were selected, ensuring the identification of meaningful associations.

The comparison of the ICARs showed conditional gene relationships shared among conditions or condition-specific for both Bone and Skin tissues. Instead of the gene expression ranges, we focused on identifying clusters of closely associated genes in the rules, using functions *as_data_frame, cluster_walktrap* and *membership* from the *igraph* package (version 2.1.1) (Csardi and Nepusz, 2006).

### 3.1.4 Baseline Analysis: Differential Gene Expression (DGE)

The traditional DGE analysis was performed on the filtered TCMs obtained after removing the Untreated condition (section 3.1.3.1), serving as a reference for evaluating the effectiveness of MettleRNASeq and as a means to assess the performance and limitations of DGE analysis when applied to these complex datasets. We selected three of the most popular and well-performing tools used for DGE analysis (Liu et al., 2022b; Corchete et al., 2020; Jiang et al., 2024): DESeq2 (version 1.44.0) (Love et al., 2014), edgeR (version 4.2.2) (Robinson et al., 2010), and limma-voom (version 3.60.6) (Ritchie et al., 2015). As with the MettleRNASeq approach, low-expression genes were filtered out, retaining only those with expression values *≥* 10 across all four replicates *per* condition, followed by normalization using the TMM method in all three DGE analyses. A less stringent adjusted p-value cutoff of 0.05 was applied to enable a fair chance of identifying differentially expressed genes. Multiple testing correction was performed using the Benjamini-Hochberg (BH) method.

For DESeq2 (v1.44.0), differential analysis was performed using the *DESeq* and *results* functions. For edgeR (v4.2.2), low-count genes were filtered with *filterByExpr*, normalized with *normLibSizes*, and analyzed using the *exactTest* function. For limma-voom (v3.60.6), the same filtering and TMM normalization functions of edgeR were applied, followed by the voom transformation with *voom*. Differential analysis was then run using *lmFit* and *eBayes* functions.

### 3.1.5 Biological Analysis

We conducted biological analyses to evaluate the relevance of key genes selected by both approaches, DGE and MettleRNASeq, for radiotherapy. Additionally, results from both tissues were examined to assess the efficacy and robustness of the MettleRNASeq approach.

Biological pathways associated with the identified genes were discovered using the Kyoto Encyclopedia of Genes and Genomes (KEGG) pathways (Kanehisa and Goto, 2000) via the *clusterProfiler* R package (version 4.12.6) (Xu et al., 2024). Gene identifiers were converted from ENSEMBL to ENTREZID using the *bitr* function set for the organism *Mus musculus* (“org.Mm.eg.db”). Enriched KEGG pathways were identified with the *enrichKEGG* function, applying a 0.05 adjusted p-value cutoff for pathway selection.

## 4. Results and Discussion

### 4.1 Datasets Analysis

Figure 3 presents the outcomes of the data variability and sample similarity analysis. All three conditions showed high variability, even among replicates, which, combined with the limited biological replicates *per* condition, further complicates the RNA-Seq analysis. Moreover, the Untreated condition contributed the most to the data variability. In contrast, the radiotherapy conditions (FLASH and Standard) showed higher similarity, highlighting the necessity to filter the Untreated condition out to obtain treatment-focused results.

**Figure 3.**
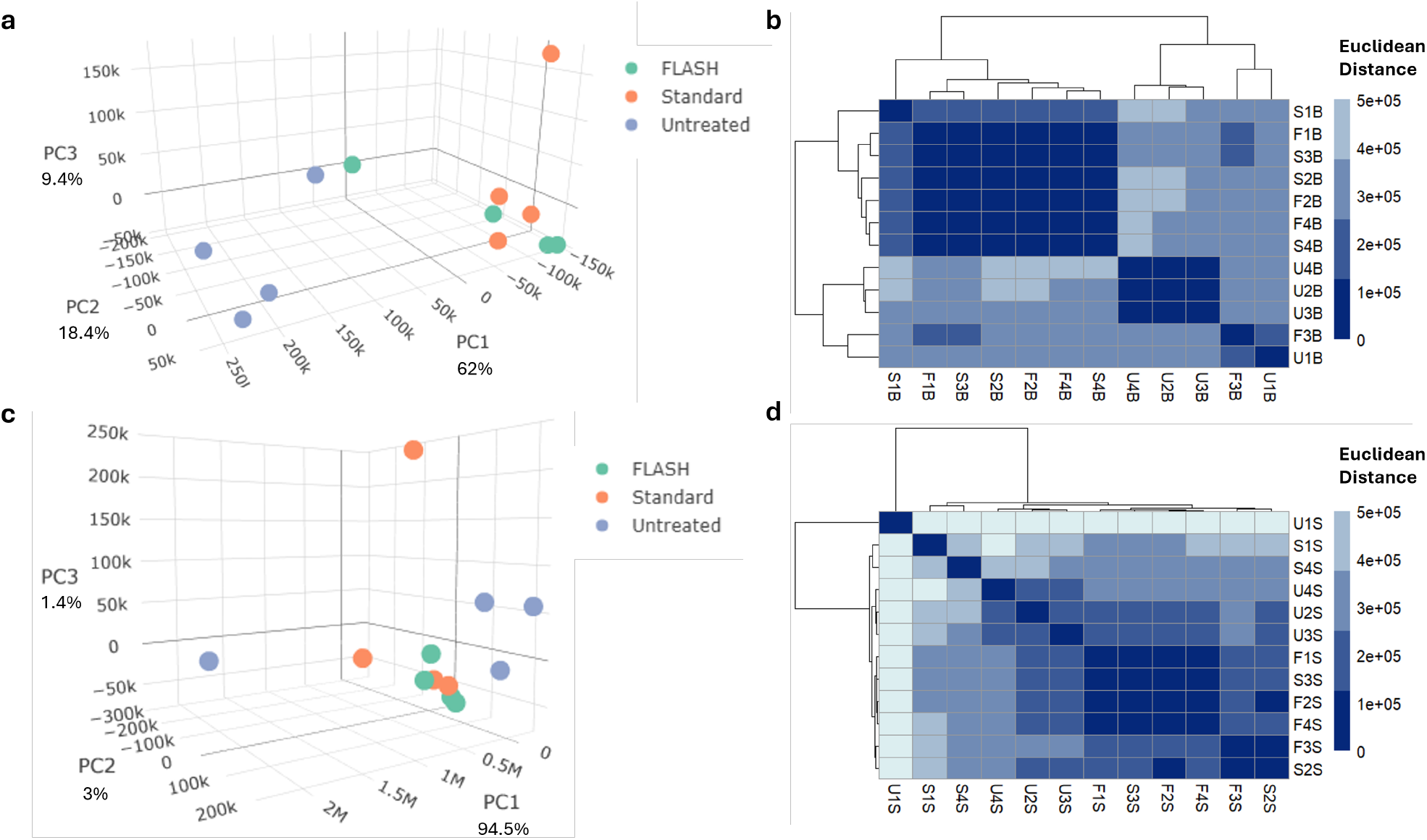
Datasets analysis. Data variability is shown in the PCAs for Bone (a) and Skin (c) tissues, in which each condition is represented by a colored dot. Heatmaps presenting the Euclidean distance illustrate sample similarity for Bone (b) and Skin (d). The row/column names identify the sample (1 to 4) and condition group (F, S, or U) from that tissue (Bone or Skin).

The Skin tissue exhibited greater sample distances (Figure 3 d), possibly reflecting tissue-specific radiation responses. As a more proliferative and radiosensitive tissue, Skin may display greater radiation-induced variability compared to Bone (Macian, 2006).

### 4.2 Traditional DGE Analysis Evaluation

The results of traditional DGE analysis using DESeq2, edgeR, and Limma-voom are presented in Figure 4.Despite the adoption of a less stringent adjusted p-value cutoff (0.05), not all DGE tools found differentially expressed genes between the treatment conditions (Figure 4 a). Notably, the Skin tissue detected nearly no DEGs, possibly due to the high variability presented by this tissue. Additionally, since both tissues underwent the same radiotherapy treatment, some overlapping DEGs between the tissues were expected, yet this was not observed with the DGE approach, as illustrated in Figure 4 b.

**Figure 4.**
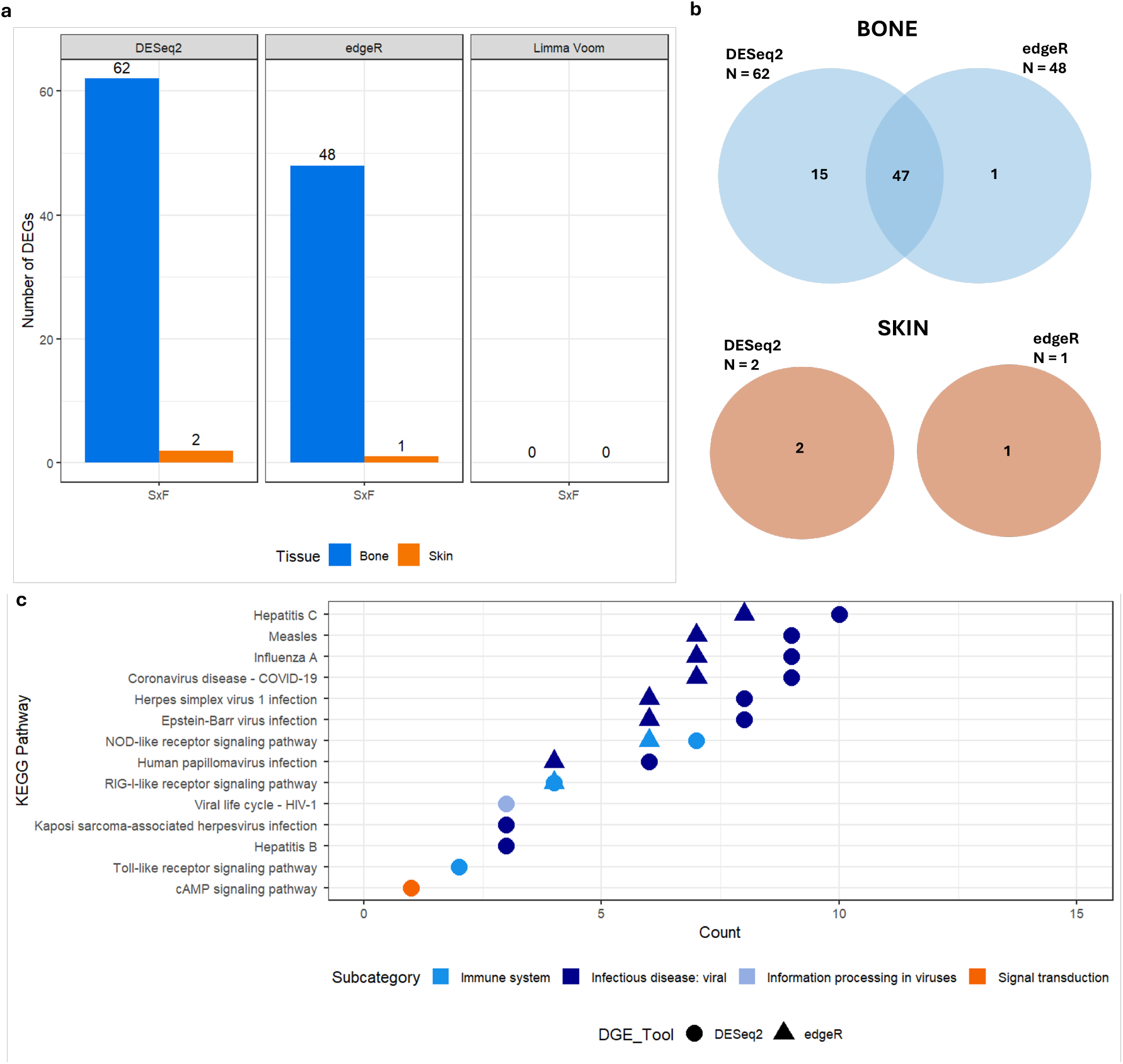
Baseline Differential Gene Expression (DGE) approach. (a) shows the number of differentially expressed genes (DEGs) found between the conditions SxF (Standard x FLASH) using DESeq2, edgeR, and Limma-voom tools (padj < 0.05) for the Bone and Skin tissues. (b) displays any intersections that occurred between the DEGs obtained by each tissue and DGE tool. (c) presents the enriched KEGG pathways in Bone (blue tones) and Skin (orange).

The Skin tissue showed only one KEGG pathway enriched (Figure 4c, in orange): the cyclic Adenosine Monophosphate (cAMP) signaling pathway. This pathway is associated with cellular radiosensitivity via the regulation of DNA repair and apoptosis (Cho et al., 2014), suggesting a link between cAMP signaling and the radiation response in Skin tissue, which is known to be more radiosensitive than Bone (Macian, 2006).

Enriched KEGG pathways in the Bone tissue include the *toll-like receptors (TLRs), NOD-like receptors (NLRs), and RIG-I-like receptors (RLRs) signaling pathways*. Those are all pattern recognition receptors (PRRs) that can trigger immune responses when sensing pathogen-associated molecular patterns (PAMPs) or danger-associated molecular patterns (DAMPs) induced by virus infection or radiation (Stoecklein et al., 2015; Spiotto et al., 2016; McGee et al., 2021). However, the fact that Bone’s enriched pathways are mostly virus-related may suggest that these PRR signaling pathways are more associated with viral infection responses within this dataset. The lack of pathways connected to radiation-specific responses may reflect the limitations of DGE analysis in this intricate data. Additionally, the absence of intersecting genes between tissues, the limited detection of DEGs across tools, and the overall challenges of statistical analysis in complex datasets further emphasize the need for an alternative strategy, such as MettleRNASeq.

### 4.3 MettleRNASeq Approach

#### 4.3.1 Classification

Resamples with the highest test accuracies (TEMs), presenting at least one TEM ≥ 70%, are shown in Figure 5. Both tissues showed variation in classification performance depending on the test sample used: Bone obtained higher accuracies with samples 2 and 4, while Skin yielded better results with samples 4 and 1. This influenced the median test accuracies, reaching up to 70% for Bone and 72% for Skin, despite some resamples hitting 90% depending on the sample used for testing.

A performance trend is also observed in Figure 5: as *S*_*r*_ increases, median test accuracies decrease, with the best results obtained at a resample size of 10. This suggests that smaller, more relevant gene sets may improve classification performance.

**Figure 5.**
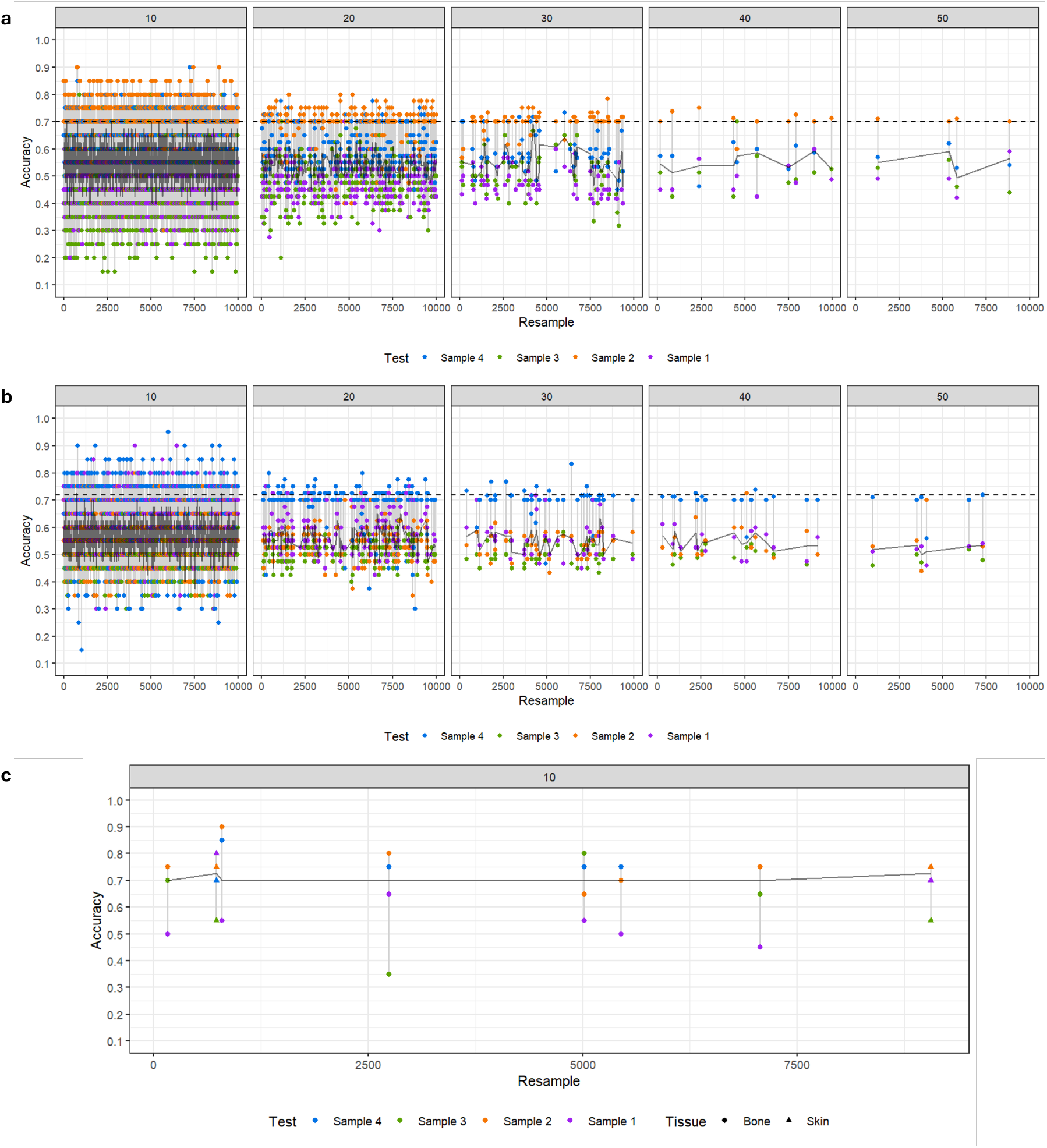
Test Accuracies for the resamples (*n* = 10.000) for Bone (a) and Skin (b). Resample sizes (*S*_*r*_) ranging from 10 to 50 are shown in each facet. Each color dot represents a test accuracy for that resample obtained by using different test samples (from 1 to 4). The solid grey line presents the median test accuracies obtained by each resample. The dashed black line highlights the median test accuracy cutoff of 70% (a) and 72% (b). In (c) the resamples selected are shown for Bone (circle) and Skin (triangle).

As shown in Figure 5 c, the resamples with the highest median test accuracies were selected (six for Bone and two for Skin) reaching 70% and 72% accuracy, respectively. From these resamples, 50 and 18 genes were identified as the best classifiers for Bone and Skin, respectively, and used for following MettleRNASeq analyses.

#### 4.3.2 MettleRNASeq’s Classification Evaluation

The genes selected after classification are presented in Figure 6 a, while those associated with the most relevant findings discussed in this section are listed in Table 1, along with their known biological roles.

**Table 1.** Genes related to the most relevant results discussed and their biological roles.

| Gene | Involved in |
| --- | --- |
| Hsp90ab1 | Maintaining protein stability under cellular stress, promoting tumor survival, proliferation, invasion, and migration (Haase and Fitze, 2016) |
| Scd1 | Regulation of key cancer-related pathways, including ferroptosis, autophagy, and immune modulation, influencing cell survival and death mechanisms (Sen et al., 2023) |
| Igha | Immune response (Dechant and Valerius, 2001) |
| Ly6c2 | Immune response (Upadhyay, 2019) |
| Hectd4 | Cell proliferation, migration, and invasion (Wan et al., 2020) |
| Tubb5 | Composition of microtubules, integral to the cytoskeleton (Seetharaman and Etienne-Manneville, 2019; Madrigal et al., 2019) |
| B2m | Tumor progression and immune regulation. Presenting tumor antigens to T cells and contributing to radiotherapy-induced immune responses (Wang et al., 2021) |
| Itm2b | Regulating apoptosis (Fleischer et al., 2002) |
| Ctss | Promoting radioresistance through suppressing BRCA1-mediated apoptosis of tumor cells (Choi et al., 2024) |
| H2-Aa | Tumor progression by regulating gene transcription and DNA damage repair (Yin et al., 2024) |
| Cybb | Ferroptosis. Shown to promote ferroptotic cancer cell death (Liu et al., 2022a) |
| Ctsk | Bone remodeling and regulation of tumor growth, and metastasis (Wu et al., 2022) |
| Serpinh1 | Cell proliferation and invasion. A cancer biomarker (Xia et al., 2024) |
| Lcem1 | Immune responses (Kang et al., 2022) |
| Anxa8 | Multiple cancer cellular processes including proliferation, metastasis, and inflammation (Zhou et al., 2021) |
| Wnk1 | Cell proliferation (with anti-apoptotic and pro-survival functions), invasion, migration, and adhesion (Jung et al., 2022) |
| Dst | Tumor suppression, cell proliferation and cytoskeleton integrity (Yu et al., 2024; Yoshioka et al., 2022) |
| Krt79 | Protection of epithelial cells from damage or stress, being a structural constituent of skin epidermis (Kim et al., 2023) |
| Eef2 | Tissue repair by regulating cellular death, autophagy, apoptosis, proliferation, and migration (Chen et al., 2023) |
| mt-Cytb | Mitochondrial electron transport chain. Related to ROS generation, playing a central role in radiation-induced stress, and mitochondrial death pathway (Lyu et al., 2024; Zhao et al., 2009; Wang et al., 2023). |
| Ly6g6c | Cancer progression and immune response (Upadhyay, 2019; Kanwal et al., 2024) |

**Figure 6.**
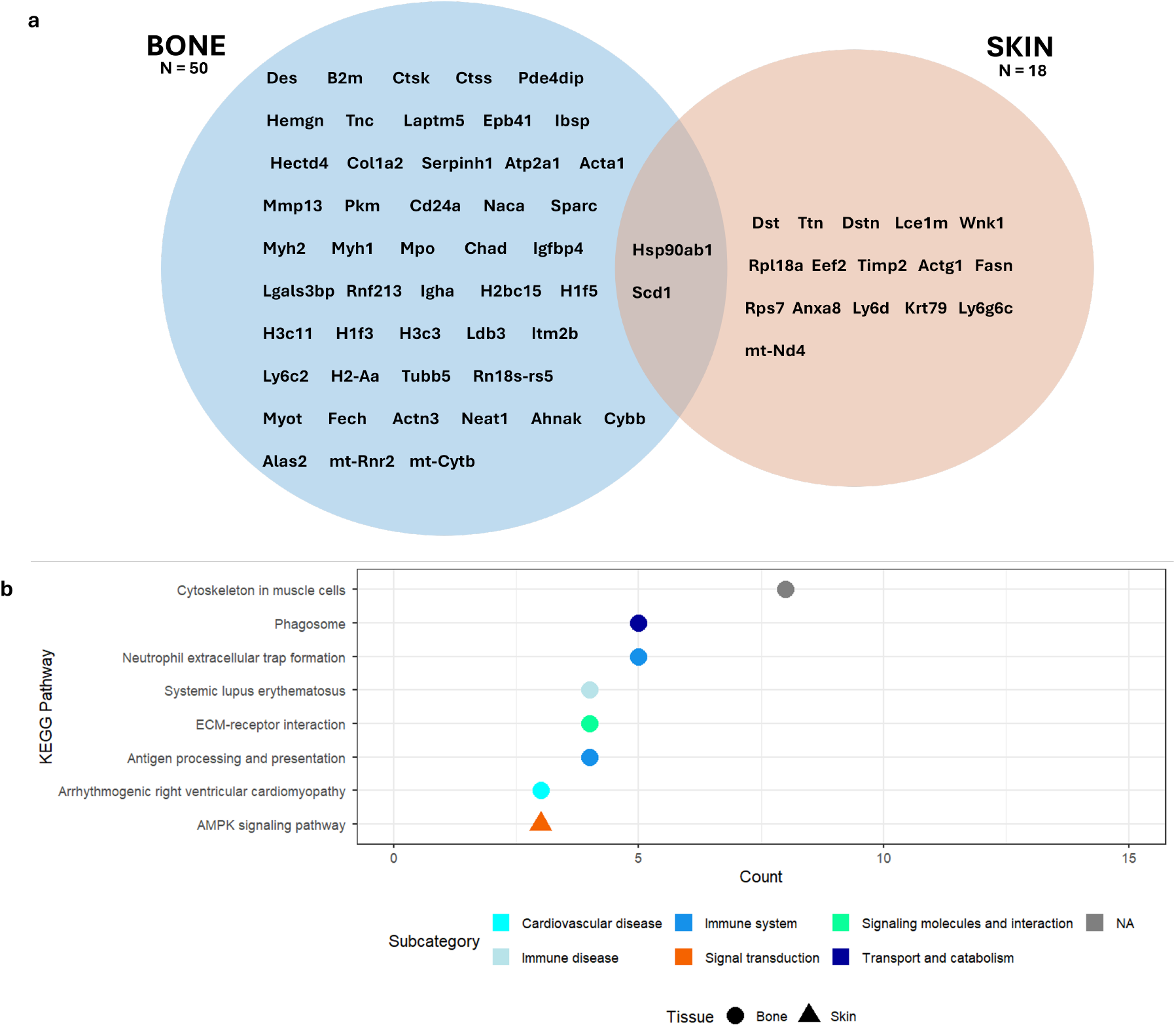
Biological analysis of MettleRNASeq. (a) shows the genes selected after classification for Bone and Skin, highlighting both unique genes and genes in common in both tissues. (b) presents the enriched KEGG pathways in the Bone (circle) and Skin (triangle) tissues.

Two genes were identified in both irradiated tissues: Hsp90ab1 and Scd1, both playing a key role in radiotherapy-induced responses by regulating cell proliferation and death, processes that are also critical for tissue regeneration following radiation-induced damage (Guerin et al., 2021).

Enriched KEGG pathways analysis highlighted diverse radiotherapy-related pathways, such as: *Cytoskeleton in muscle cells, Phagosome, Neutrophil Extracelular Trap formation, ECM-receptor interaction, Antigen processing and presentation* and *AMPK signaling pathway*. Radiation is known to alter *cytoskeleton* components (actin filaments, intermediate filaments, and microtubules), affecting adhesion, migration, and treatment response (Toffali et al., 2023); increase *phagocytic* activity of macrophages (Lecoultre et al., 2024; Lee et al., 2020); induce *extracellular matrix (ECM)* disorder promoting malignancy and therapy resistance (Yan et al., 2024); promote immune responses, being highly connected with *Neutrophil extracellular trap (NET) formation* and *Antigen processing and presentation* (Shinde-Jadhav et al., 2021; Tailor et al., 2022; Pishesha et al., 2022); and activate stress responses regulated by *AMP-activated protein kinase (AMPK)* (Zannella et al., 2011). These enriched pathways reinforce the relevance of the identified genes in the context of radiotherapy, supporting the robustness of MettleRNASeq when applied to complex datasets. Such results enable further research to deepen biological insights, including comparisons between FLASH and Standard radiotherapy, as well as the exploration of gene relationships.

When comparing the genes selected by DGE and MettleRNASeq, no overlaps were found in Skin, while only three genes overlapped in Bone: Igha, Ly6c2, and Rnf213, linked to processes like immune response, angiogenesis and cell proliferation, migration, and invasion (Lin et al., 2024; Ye et al., 2023; Zhang et al., 2023; Chi-ablaem et al., 2024; Dechant and Valerius, 2001; Upadhyay, 2019). Notably, in Skin both approaches highlighted related pathways (AMPK and cAMP), connected through the Ataxia Telangiectasia Mutated (ATM) protein, and involved in radiation-induced DNA damage responses and cell death (Li et al., 2017; Cho et al., 2014).

#### 4.3.3 Correlation Analysis

Correlation analysis generated ACC and ICC results, which are illustrated in Figures 7, 8, and 9. The ACC results revealed strong positive and negative correlations (| correlation | ≥ 0.7) between gene expressions in FLASH and Standard conditions, possibly reflecting key biological processes involved in the radiotherapy response, either being crucial for both treatments (positive ACCs) or condition-specific (negative ACCs). In both tissues, the majority of strong ACCs were positive: in Bone (Figure 7 a) notable genes include Hectd4 and Tubb5 (Table 1), linked to cell adhesion and migration, affecting treatment resistance and cellular adaptation (Seetharaman and Etienne-Manneville, 2019; Madrigal et al., 2019); in Skin (Figure 7 b), genes Hsp90ab1, Anxa8, Wnk1, and Dst can be emphasized, involved in essential processes like cell proliferation. In contrast, strong negative ACCs revealed greater variance in gene expression between conditions, encompassing genes B2m, Itm2b, and Ctss in Bone tissue (Figure 7 a), linked to immune responses and cell death, which are processes identified by several studies to be differentially regulated by FLASH and Standard radiotherapy (Chow and Ruda, 2024; Alhaddad et al., 2024; Allen et al., 2020). In Skin, gene Lcem1 (also known as KKLC1) was the only one to exhibit a strong negative ACC, being associated with triggering immune responses, a recognized mechanism of the FLASH effect for an enhanced repair of the normal tissue injured by radiation (Kang et al., 2022; Alhad-dad et al., 2024).

**Figure 7.**
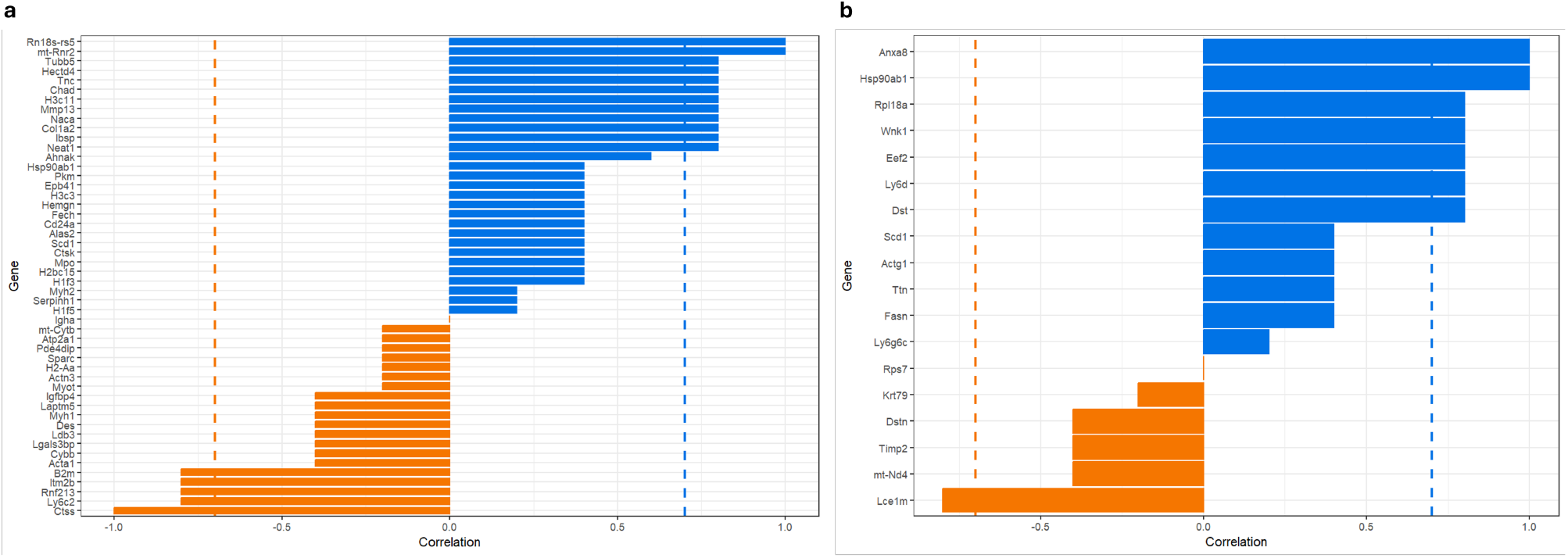
Across-Condition Spearman Correlations (ACC) for each tissue. The correlation showing the relationship between the two treatment conditions by gene is shown for the Bone in (a) and Skin in (b). Positive and negative correlations are presented in blue and orange, respectively. The dashed line highlights the start of stronger correlations (|correlation| *≥* 0.7).

**Figure 8.**
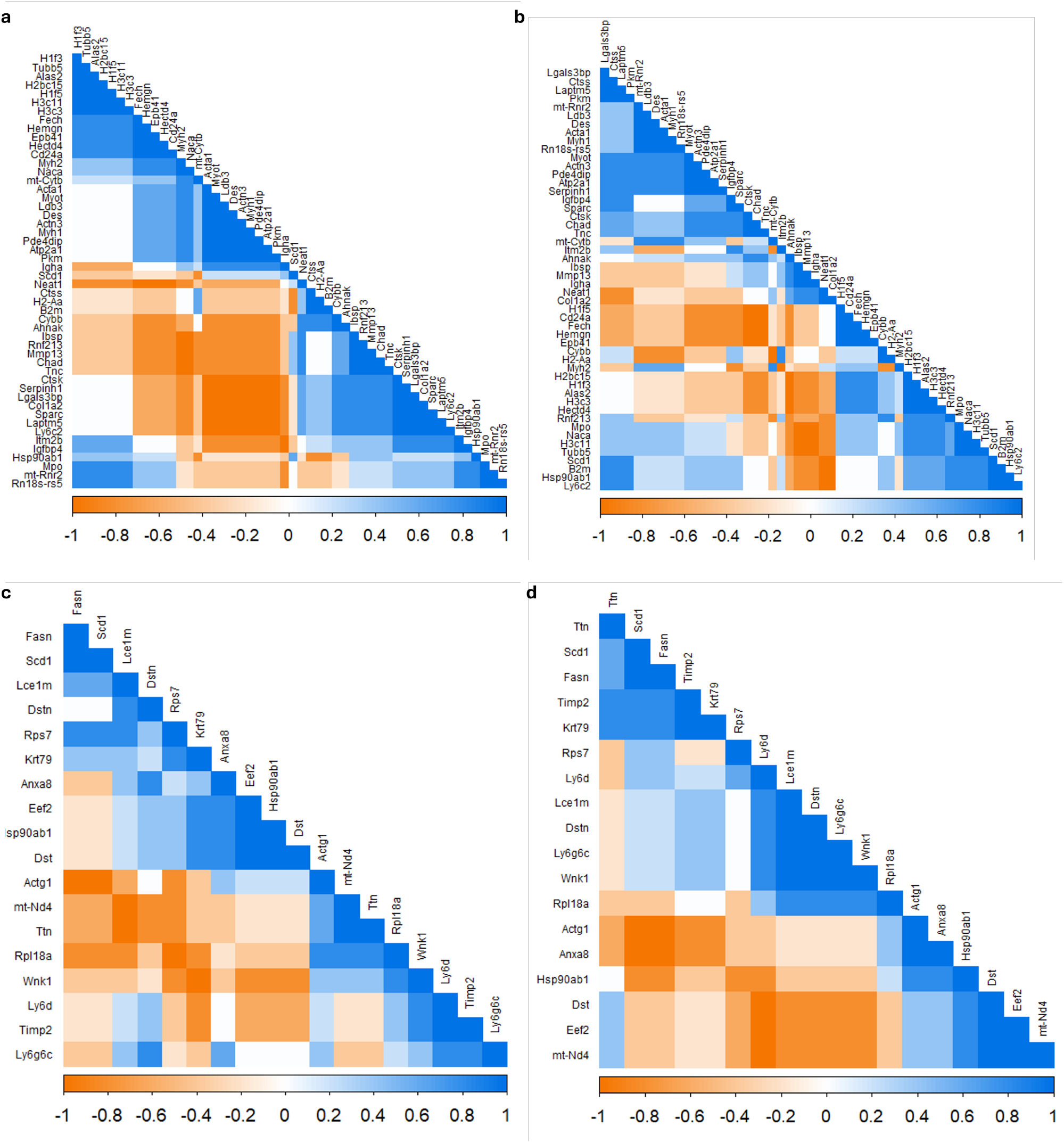
Intra-Condition Spearman Correlations (ICC) for each tissue. Correlation coefficients between gene pairs in Bone tissue are shown in (a) for FLASH and (b) for Standard. In Skin tissue, the correlations are displayed in (c) for FLASH and (d) for Standard. Positive and negative correlations are presented in blue and orange, respectively, and the stronger the color the stronger the correlation between the genes.

**Figure 9.**
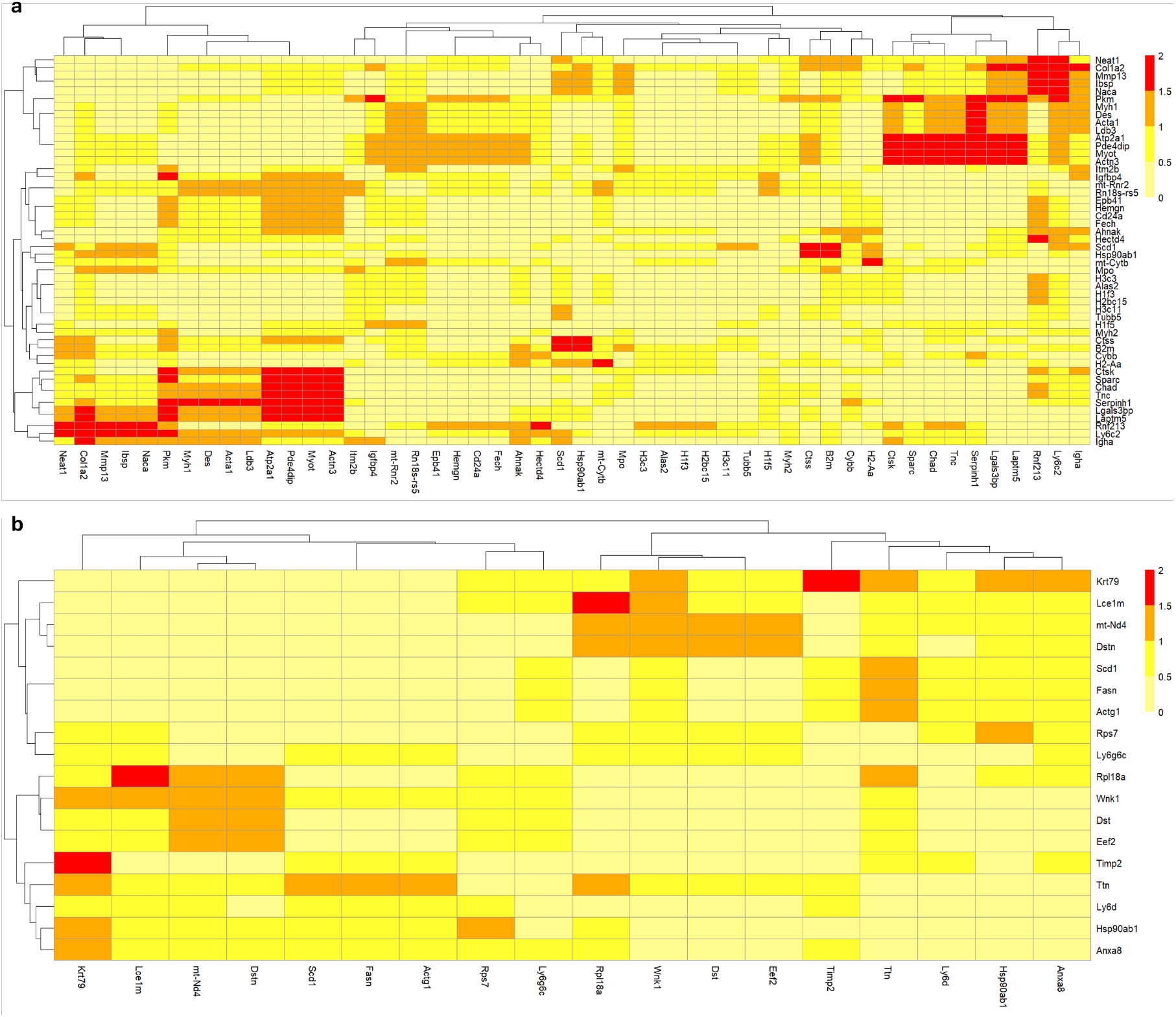
Absolute correlation differences between Intra-Condition Correlations (ICCs) for each tissue. (a) Bone and (b) Skin. Stronger colors indicate greater differences in gene correlations between conditions within each tissue.

Additionally, shared or tissue-specific responses can also be observed in Figure 7. For instance, the Scd1 gene maintained a weak correlation in both tissues; however, Hsp90ab1 showed varied ACCs, exhibiting a weak ACC in Bone but a strong positive ACC in Skin. Given Hsp90ab1’s essential role following radiation-induced stress, this variation may reflect tissue-specific responses to radiotherapy, potentially linked to differences in radiosensitivity or radioresistance.

Interestingly, even genes with weak ACCs, such as Scd1, may still offer relevant insights, particularly through their relationships rather than solely through changes in their own expression. For instance, despite its ACC results, Scd1 showed strong negative ICCs with genes like H2-Aa, mt-Cytb, B2m, and Ctss in Bone (FLASH, Figure 8 a). These genes are all involved in cell death and damage repair, processes closely related to radiotherapy response (Amundson, 2022; Chen and Kuo, 2017; Lu et al., 2022).

The ICC results presented in Figure 8 revealed several gene pairs with strong correlations, suggesting potential functional or regulatory relationships. For example, in Bone (FLASH, Figure 8 a), genes Tubb5 and Hectd4 showed strong positive correlations, while Serpinh1 and Ctsk displayed strong negative correlations. However, the ICCs can change or persist depending on the treatment; shared or condition-specific regulations of gene relationships were explored in Figure 9, in which some of the greatest correlation changes between treatments in Bone and Skin included genes Serpinh1 and Krt79, respectively, possibly explaining the different radiotherapy responses at play. For instance, since Krt79 is related to the protection of epithelial cells and the Skin tissue is more sensitive to radiation, the notable difference in this gene’s expression between the conditions could be linked to the different levels of damage and repair caused by the radiotherapies (Chow and Ruda, 2024). In contrast, smaller differences in gene correlations between conditions may also be of interest, as they could indicate essential or non-essential processes shared between both radiotherapy treatments: strong correlations that remain largely unchanged involve genes Hectd4 and Ly6g6c in Bone and Skin, respectively, being related to cell proliferation, migration, invasion, and immune response; alternatively, weaker correlations that persist across conditions may represent baseline interactions unrelated to radiation exposure.

#### 4.3.4 Association Rule Mining

Conditional gene relationships, highlighting shared and condition-specific gene relations, are shown in Figure 10. Interestingly, Scd1 and Hsp90ab1, the only genes identified in both tissues as seen in Figure 5, appeared in shared rules across conditions for Bone and Skin (Figure 10 a and d), reinforcing their relevance in radiotherapy response through their roles in cell death, proliferation, and tissue regeneration (Guerin et al., 2021). Yet, despite their relevance, they showed distinct gene associations in each tissue: while in Bone (a), Scd1 and Hsp90ab1 are part of the same rule, in Skin (d) they are associated with different genes, e.g., Hsp90ab1 appears in the same cluster as Dst and Eef2.

**Figure 10.**
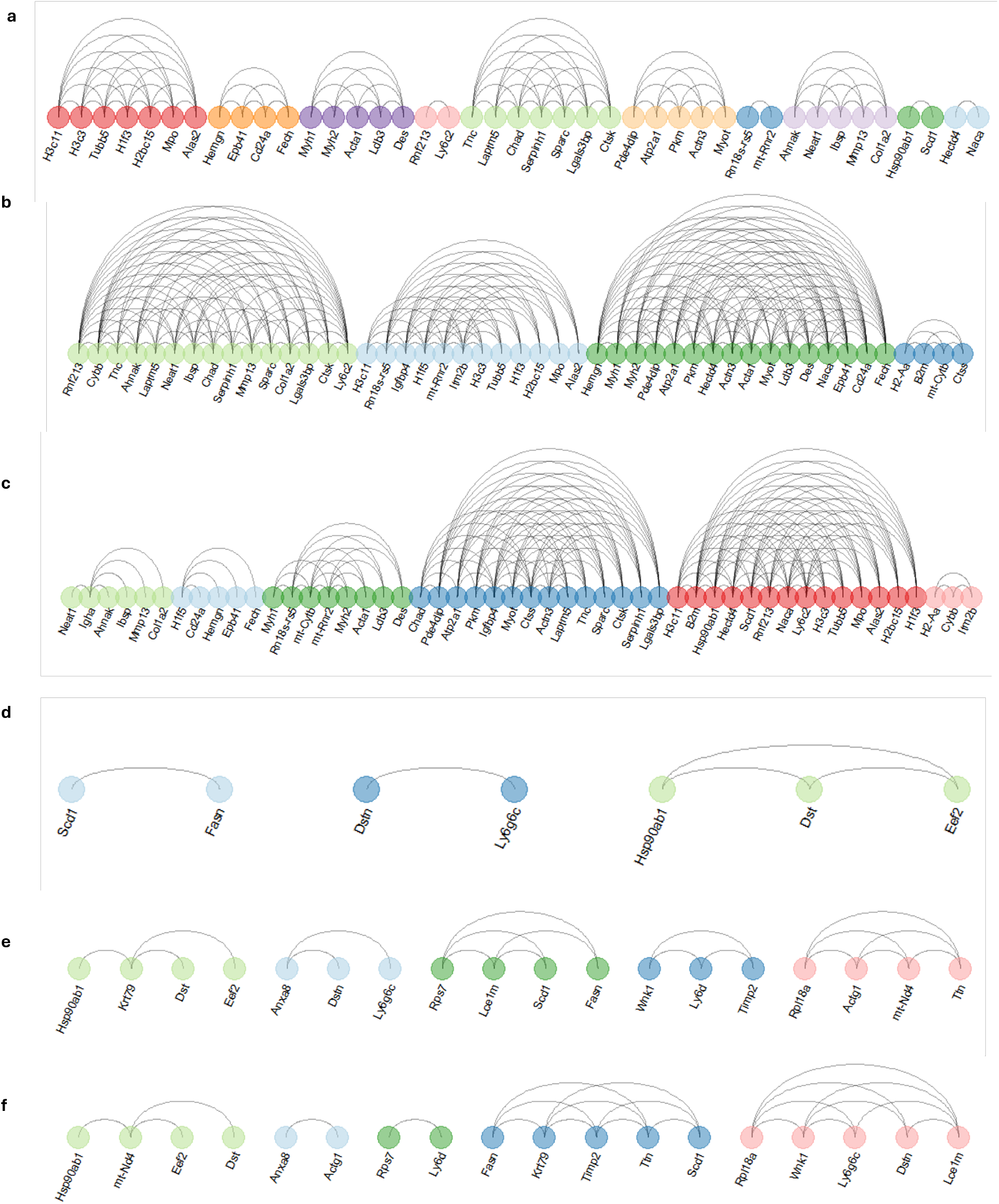
Rules of Association found between genes for each tissue. Colored dots indicate gene clusters within the same rule. Shared and condition-specific (FLASH and Standard) rules are shown for Bone in (a–c), and for Skin in (d–f), respectively.

Condition-specific gene relations were also identified: in the Bone tissue, FLASH radiotherapy presented a gene cluster between H2-Aa and genes B2m, mt-Cytb, and Ctss; in contrast, in Standard radiotherapy, the H2-Aa gene formed a cluster with Cybb and Itm2b. Although both clusters involve processes related to immune response, oxidative stress, cell death, and damage repair, the formation of distinct clusters may highlight deeper underlying mechanisms triggered by each radiotherapy, leading to differential treatment outcomes.

## 5. Conclusion

Traditional DGE approaches are widely used to identify differentially expressed genes, but they have inherent limitations, particularly in complex RNA-Seq datasets. These approaches focus on statistical significance, making them susceptible to false positives and negatives in datasets with high variability, limited biological replicates *per* condition, and intricate biological mechanisms. Additionally, by overlooking gene relationships, traditional DGE may miss meaningful interactions that drive biological phenomena.

MettleRNASeq was designed to overcome these challenges by moving beyond statistical significance. By identifying key genes that accurately distinguish conditions and highlighting gene relationships, it uncovers condition-specific and shared biological processes that traditional DGE approaches might miss. MettleRNASeq has been validated, demonstrating its effectiveness and consistency in complex biological systems. This suggests that our approach can serve as a valuable tool for analyzing intricate RNA-Seq data, offering a thorough study of gene expression.

## Data Availability

Datasets can be retrieved from the Gene Expression Omnibus (GEO) database at NCBI (Accession GSE173944). MettleRNASeq R package is available on GitHub at: https://github.com/SamellaSalles/MettleRNASeq The supplementary data of this study are available from the corresponding author upon reasonable request.

## Competing interests

No competing interest is declared.

## Author contributions statement

S.S. collected and prepared the data. S.S., O.B., L.C., and K.O. conceived the experiment. S.S. and O.B. conducted the experiment. S.S., O.B., L.C., and K.O. analyzed the results. S.S. wrote the manuscript. S.S., O.B., L.C., and K.O. reviewed the manuscript.

## Acknowledgments

The authors thank the National Laboratory for Scientific Computing (LNCC, Brazil) for providing support and access to the Santos Dumont (SDumont) Supercomputer. We also acknowledge the Coordination for the Improvement of Higher Education Personnel (CAPES) for funding this research. Special thanks to Rebecca Salles for the valuable discussions and feedback.

## Notes

https://github.com/SamellaSalles/MettleRNASeq/

### Competing Interest Statement

The authors have declared no competing interest.

## Bibliography

Akobeng, A. K. (2016). Understanding type i and type ii errors, statistical power and sample size. Acta Paediatrica, 105(6):605–609.

Alhaddad, L., Osipov, A. N., and Leonov, S. (2024). Flash radiotherapy: Benefits, mechanisms, and obstacles to its clinical application. International Journal of Molecular Sciences, 25(23):12506.

Allen, B. D., Acharya, M. M., Montay-Gruel, P., Jorge, P. G., Bailat, C., Petit, B., Vozenin, M.-C., and Limoli, C. (2020). Maintenance of tight junction integrity in the absence of vascular dilation in the brain of mice exposed to ultra-high-dose-rate flash irradiation. Radiation research, 194(6):625–635.

Amundson, S. A. (2022). The transcriptomic revolution and radiation biology. International journal of radiation biology, 98(3):428–438.

Anders, S., Pyl, P. T., and Huber, W. (2015). Htseq—a python framework to work with high-throughput sequencing data. bioinformatics, 31(2):166–169.

Andrews, S. et al. Fastqc: a quality control tool for high throughput sequence data, (2010).

Arowolo, M. O., Adebiyi, M. O., Aremu, C., and Adebiyi, A. A. (2021). A survey of dimension reduction and classification methods for rna-seq data on malaria vector. Journal of Big Data, 8:1–17.

Bishop, C. M. and Nasrabadi, N. M. Pattern recognition and machine learning, volume 4. Springer, (2006).

Bradshaw, T. J., Huemann, Z., Hu, J., and Rahmim, A. (2023). A guide to cross-validation for artificial intelligence in medical imaging. Radiology: Artificial Intelligence, 5(4):e220232.

Breiman, L. (2001). Random forests. Machine learning, 45:5–32.

Cassidy, J., Bissett, D., OBE, R. A. S., Payne, M., and Morris-Stiff, G. Oxford handbook of oncology. OUP Oxford, (2015).

Chavan-Gautam, P., Shah, T., Joshi, K., Patwardhan, B., and Chaguturu, R. (2017). Transcriptomics and epigenomics. Innovative Approaches in Drug Discovery: Ethnopharmacology, Systems Biology and Holistic Targeting. Elsevier Inc. 10.1016/B978-0-12-801814-9.00008-8.

Chen, H. H. and Kuo, M. T. (2017). Improving radiotherapy in cancer treatment: Promises and challenges. Oncotarget, 8(37):62742.

Chen, J., Wang, J., Wu, X., Simon, N., Svensson, C. I., Yuan, J., Hart, D. A., Ahmed, A. S., and Ackermann, P. W. (2023). eef2 improves dense connective tissue repair and healing outcome by regulating cellular death, autophagy, apoptosis, proliferation and migration. Cellular and Molecular Life Sciences, 80(5):128.

Chen, J. W. and Dhahbi, J. (2021). Lung adenocarcinoma and lung squamous cell carcinoma cancer classification, biomarker identification, and gene expression analysis using overlapping feature selection methods. Scientific reports, 11(1):13323.

Chen, S.-C., Tsai, T.-H., Chung, C.-H., and Li, W.-H. (2015). Dynamic association rules for gene expression data analysis. BMC genomics, 16:1–20.

Chen, Y., Chen, S., Wu, Z., Cheng, Q., and Ji, D. (2024). Hypoxia-related lncrna correlates with prognosis and immune microenvironment in uveal melanoma. Cancer Cell International, 24(1):336.

Cheng, Y., Xu, S.-M., Santucci, K., Lindner, G., and Janitz, M. (2024). Machine learning and related approaches in transcriptomics. Biochemical and Biophysical Research Communications, page 150225.

Chiablaem, K., Jinawath, A., Nuanpirom, J., Arora, J. K., Nasaree, S., Thanomchard, T., Singhto, N., Chittavanich, P., Suktitipat, B., Charoensawan, V., et al. (2024). Identification of rnf213 as a potential suppressor of local invasion in intrahepatic cholangiocarcinoma. Laboratory Investigation, page 102074.

Ching, T., Huang, S., and Garmire, L. X. (2014). Power analysis and sample size estimation for rna-seq differential expression. Rna, 20(11):1684–1696.

Cho, E.-A., Kim, E.-J., Kwak, S.-J., and Juhnn, Y.-S. (2014). camp signaling inhibits radiation-induced atm phosphorylation leading to the augmentation of apoptosis in human lung cancer cells. Molecular cancer, 13:1–15.

Choi, E., Jeon, K.-H., Lee, H., Mun, G.-I., Kim, J.-A., Shin, J.-H., Kwon, Y., Na, Y., and Lee, Y.-S. (2024). Radiosensitizing effect of a novel ctss inhibitor by enhancing brca1 protein stability in triple-negative breast cancer cells. Cancer Science.

Chow, J. C. and Ruda, H. E. (2024). Mechanisms of action in flash radiotherapy: A comprehensive review of physicochemical and biological processes on cancerous and normal cells. Cells, 13(10):835.

Corchete, L. A., Rojas, E. A., Alonso-López, D., De Las Rivas, J., Gutiérrez, N. C., and Burguillo, F. J. (2020). Systematic comparison and assessment of rna-seq procedures for gene expression quantitative analysis. Scientific reports, 10(1):19737.

Csardi, G. and Nepusz, T. (2006). The igraph software. Complex syst, 1695:1–9.

Dechant, M. and Valerius, T. (2001). Iga antibodies for cancer therapy. Critical reviews in oncology/hematology, 39(1-2):69–77.

Dobin, A., Davis, C. A., Schlesinger, F., Drenkow, J., Zaleski, C., Jha, S., Batut, P., Chaisson, M., and Gingeras, T. R. (2013). Star: ultrafast universal rna-seq aligner. Bioinformatics, 29(1):15–21.

Du, C., Wang, C., Liu, Z., Xin, W., Zhang, Q., Ali, A., Zeng, X., Li, Z., and Ma, C. (2024). Machine learning algorithms integrate bulk and single-cell rna data to unveil oxidative stress following intracerebral hemorrhage. International Immunopharmacology, 137:112449.

Essam, F., El, H., and Ali, S. R. H. (2022). A comparison of the pearson, spearman rank and kendall tau correlation coefficients using quantitative variables. Asian J. Probab. Stat, pages 36–48.

Ewels, P., Magnusson, M., Lundin, S., and Käller, M. (2016). Multiqc: summarize analysis results for multiple tools and samples in a single report. Bioinformatics, 32(19):3047– 3048.

Fleischer, A., Ayllon, V., and Rebollo, A. (2002). Itm2bs regulates apoptosis by inducing loss of mitochondrial membrane potential. European journal of immunology, 32(12):3498– 3505.

Fonseca, N. A., Marioni, J., and Brazma, A. (2014). Rna-seq gene profiling-a systematic empirical comparison. Plos one, 9(9):e107026.

Frankish, A., Carbonell-Sala, S., Diekhans, M., Jungreis, I., Loveland, J. E., Mudge, J. M., Sisu, C., Wright, J. C., Arnan, C., Barnes, I., et al. (2023). Gencode: reference annotation for the human and mouse genomes in 2023. Nucleic acids research, 51(D1):D942–D949.

Gakii, C. and Rimiru, R. (2021). Identification of cancer related genes using feature selection and association rule mining. Informatics in Medicine Unlocked, 24:100595.

Guerin, D. J., Kha, C. X., and Tseng, K. A.-S. (2021). From cell death to regeneration: rebuilding after injury. Frontiers in Cell and Developmental Biology, 9:655048.

Haase, M. and Fitze, G. (2016). Hsp90ab1: Helping the good and the bad. Gene, 575(2):171–186.

Hahsler, M., Gruen, B., and Hornik, K. (2005). arules – A computational environment for mining association rules and frequent item sets. Journal of Statistical Software, 14(15):1–25. doi: 10.18637/jss.v014.i15.

Hegland, M. (2007). The apriori algorithm–a tutorial. Mathematics and computation in imaging science and information processing, pages 209–262.

Jabeen, A., Ahmad, N., and Raza, K. Machine learning-based state-of-the-art methods for the classification of rna-seq data. In Classification in BioApps: Automation of Decision Making, pages 133–172. Springer, (2017).

Jiang, G., Zheng, J.-Y., Ren, S.-N., Yin, W., Xia, X., Li, Y., and Wang, H.-L. (2024). A comprehensive workflow for optimizing rna-seq data analysis. BMC genomics, 25(1):631.

Jung, J.-U., Jaykumar, A. B., and Cobb, M. H. (2022). Wnk1 in malignant behaviors: A potential target for cancer? Frontiers in Cell and Developmental Biology, 10:935318.

Kanehisa, M. and Goto, S. (2000). Kegg: kyoto encyclopedia of genes and genomes. Nucleic acids research, 28(1):27–30.

Kang, Y., Gan, Y., Jiang, Y., You, J., Huang, C., Chen, Q., Xu, X., Chen, F., and Chen, L. (2022). Cancer-testis antigen kk-lc-1 is a potential biomarker associated with immune cell infiltration in lung adenocarcinoma. BMC cancer, 22(1):834.

Kanwal, M., Smahelova, J., Ciharova, B., Johari, S. D., Nunvar, J., Olsen, M., and Smahel, M. (2024). Aspartate β-hydroxylase regulates expression of ly6 genes. Journal of Cancer, 15(5):1138.

Kim, D., Choi, I., Ha, S. K., and Gonzalez, F. J. (2023). Keratin 79 is a ppara target that is highly expressed by liver damage. Biochemical and biophysical research communications, 650:132–136.

Kitchin, R. and Lauriault, T. P. (2015). Small data in the era of big data. GeoJournal, 80:463–475.

Koch, C. M., Chiu, S. F., Akbarpour, M., Bharat, A., Ridge, K. M., Bartom, E. T., and Winter, D. R. (2018). A beginner’s guide to analysis of rna sequencing data. American journal of respiratory cell and molecular biology, 59(2):145–157.

Kolde, R. pheatmap: Pretty Heatmaps, (2019). URL https://CRAN.R-project.org/package=pheatmap. R package version 1.0.12.

Kuhn and Max. (2008). Building predictive models in r using the caret package. Journal of Statistical Software, 28(5):1–26. doi: 10.18637/jss.v028.i05.

Lakens, D. (2021). The practical alternative to the p value is the correctly used p value. Perspectives on psychological science, 16(3):639–648.

Lecoultre, M., Chliate, S., Espinoza, F. I., Tankov, S., Dutoit, V., and Walker, P. R. (2024). Radio-chemotherapy of glioblastoma cells promotes phagocytosis by macrophages in vitro. Radiotherapy and Oncology, 190:110049.

Lee, H.-J., Woo, Y., Hahn, T.-W., Jung, Y. M., and Jung, Y.-J. (2020). Formation and maturation of the phagosome: a key mechanism in innate immunity against intracellular bacterial infection. Microorganisms, 8(9):1298.

Lemoine, G. G., Scott-Boyer, M.-P., Ambroise, B., Périn, O., and Droit, A. (2021). Gwena: gene co-expression networks analysis and extended modules characterization in a single bioconductor package. BMC bioinformatics, 22(1):267.

Li, R., Luo, X., Zhu, Y., Zhao, L., Li, L., Peng, Q., Ma, M., and Gao, Y. (2017). Atm signals to ampk to promote autophagy and positively regulate dna damage in response to cadmiuminduced ros in mouse spermatocytes. Environmental Pollution, 231:1560–1568.

Lin, B., Gao, F., Yang, Y., Wu, D., Zhang, Y., Feng, G., Dai, T., and Du, X. (2021). Flash radiotherapy: history and future. Frontiers in oncology, 11:644400.

Lin, T.-C., Uchino, H., Ito, M., Yamaguchi, S., Ishi, Y., and Fujimura, M. (2024). Moyamoya syndrome after proton beam therapy in a pediatric patient with a pineal germ cell tumor and a germline polymorphism in rnf213. Child’s Nervous System, 40(11):3873–3878.

Lin, W.-Y., Tseng, M.-C., and Su, J.-H. A confidence-lift support specification for interesting associations mining. In Advances in Knowledge Discovery and Data Mining: 6th Pacific-Asia Conference, PAKDD 2002 Taipei, Taiwan, May 6–8, 2002 Proceedings 6, pages 148–158. Springer, (2002).

Liu, J., Kang, R., and Tang, D. (2022). Signaling pathways and defense mechanisms of ferroptosis. The FEBS journal, 289(22):7038–7050.

Liu, X., Zhao, J., Xue, L., Zhao, T., Ding, W., Han, Y., and Ye, H. (2022). A comparison of transcriptome analysis methods with reference genome. Bmc Genomics, 23(1):232.

Liu, Y., Yao, Y., Zhang, Y., Xu, C., Yang, T., Qu, M., Lu, B., Song, X., Pan, X., Zhou, W., et al. (2024). Identification of prognostic stemness-related genes in kidney renal papillary cell carcinoma. BMC Medical Genomics, 17(1):121.

Love, M. I., Huber, W., and Anders, S. (2014). Moderated estimation of fold change and dispersion for rna-seq data with deseq2. Genome biology, 15:1–21.

Lowe, R., Shirley, N., Bleackley, M., Dolan, S., and Shafee, T. (2017). Transcriptomics technologies. PLoS computational biology, 13(5):e1005457.

Lu, Y. and Belitskaya-Levy, I. (2015). The debate about p-values. Shanghai Archives of Psychiatry, 27(6):381–385.

Lu, Z., Zheng, X., Ding, C., Zou, Z., Liang, Y., Zhou, Y., and Li, X. (2022). Deciphering the biological effects of radiotherapy in cancer cells. Biomolecules, 12(9):1167.

Lv, Y., Lv, Y., Wang, Z., Lan, T., Feng, X., Chen, H., Zhu, J., Ma, X., Du, J., Hou, G., et al. (2022). Flash radiotherapy: A promising new method for radiotherapy. Oncology letters, 24(6):1–14.

Lyu, X., Yu, Y., Jiang, Y., Li, Z., and Qiao, Q. (2024). The role of mitochondria transfer in cancer biological behavior, the immune system and therapeutic resistance. Journal of Pharmaceutical Analysis, page 101141.

Macian, R. (2006). Biological effects of radiation. Reactors Concepts Manual, USNRC Technical Training Center.

Madrigal, I., Rabionet, R., Alvarez-Mora, M., Sanchez, A., Rodríguez-Revenga, L., Estivill, X., and Mila, M. (2019). Spectrum of clinical heterogeneity of β-tubulin tubb5 gene mutations. Gene, 695:12–17.

Martin, M. (2011). Cutadapt removes adapter sequences from high-throughput sequencing reads. EMBnet. journal, 17(1):10–12.

Matuszak, N., Suchorska, W. M., Milecki, P., Kruszyna-Mochalska, M., Misiarz, A., Pracz, J., and Malicki, J. (2022). Flash radiotherapy: an emerging approach in radiation therapy. reports of Practical Oncology and radiotherapy, 27(2):343–351.

McGee, H. M., Marciscano, A. E., Campbell, A. M., Monjazeb, A. M., Kaech, S. M., and Teijaro, J. R. (2021). Parallels between the antiviral state and the irradiated state. JNCI: Journal of the National Cancer Institute, 113(8):969–979.

Pishesha, N., Harmand, T. J., and Ploegh, H. L. (2022). A guide to antigen processing and presentation. Nature Reviews Immunology, 22(12):751–764.

Qian, J., Liu, W., Shi, Y., Zhang, M., Wu, Q., Chen, K., and Chen, W. (2022). C-cora: A cluster-based method for correlation analysis of rna-seq data. Horticulturae, 8(2):124.

R Core Team. R: A Language and Environment for Statistical Computing. R Foundation for Statistical Computing, Vienna, Austria, (2024). URL https://www.R-project.org/.

Ritchie, M. E., Phipson, B., Wu, D., Hu, Y., Law, C. W., Shi, W., and Smyth, G. K. (2015). limma powers differential expression analyses for rna-sequencing and microarray studies. Nucleic acids research, 43(7):e47–e47.

Robinson, M. D. and Oshlack, A. (2010). A scaling normalization method for differential expression analysis of rna-seq data. Genome biology, 11:1–9.

Robinson, M. D., McCarthy, D. J., and Smyth, G. K. (2010). edger: a bioconductor package for differential expression analysis of digital gene expression data. bioinformatics, 26(1):139–140.

Ruan, J., Dean, A. K., and Zhang, W. (2010). A general co-expression network-based approach to gene expression analysis: comparison and applications. BMC systems biology, 4:1–21.

Seetharaman, S. and Etienne-Manneville, S. (2019). Microtubules at focal adhesions–a double-edged sword. Journal of cell science, 132(19):jcs232843.

Sen, U., Coleman, C., and Sen, T. (2023). Stearoyl coenzyme a desaturase-1: multitasker in cancer, metabolism, and ferroptosis. Trends in cancer, 9(6):480–489.

Shinde-Jadhav, S., Mansure, J. J., Rayes, R. F., Marcq, G., Ayoub, M., Skowronski, R., Kool, R., Bourdeau, F., Brimo, F., Spicer, J., et al. (2021). Role of neutrophil extracellular traps in radiation resistance of invasive bladder cancer. Nature communications, 12(1):2776.

Sievert, C. Interactive Web-Based Data Visualization with R, plotly, and shiny. Chapman and Hall/CRC, (2020). ISBN 9781138331457. URL https://plotly-r.com.

Skerrett-Byrne, D. A., Chen, J. C., Nixon, B., and Hondermarck, H. (2023). Transcriptomics. Encyclopedia of Cell Biology (Second Edition), pages 363–371.

Spiotto, M., Fu, Y.-X., and Weichselbaum, R. R. (2016). The intersection of radiotherapy and immunotherapy: mechanisms and clinical implications. Science immunology, 1(3):eaag1266–eaag1266.

Stoecklein, V. M., Osuka, A., Ishikawa, S., Lederer, M. R., Wanke-Jellinek, L., and Lederer, J. A. (2015). Radiation exposure induces inflammasome pathway activation in immune cells. The Journal of Immunology, 194(3):1178–1189.

Su, K., Wu, Z., and Wu, H. (2020). Simulation, power evaluation and sample size recommendation for single-cell rna-seq. Bioinformatics, 36(19):4860–4868.

Swanton, C., Bernard, E., Abbosh, C., André, F., Auwerx, J., Balmain, A., Bar-Sagi, D., Bernards, R., Bullman, S., DeGregori, J., et al. (2024). Embracing cancer complexity: Hallmarks of systemic disease. Cell, 187(7):1589–1616.

Tailor, A., Estephan, H., Parker, R., Woodhouse, I., Abdulghani, M., Nicastri, A., Jones, K., Salatino, S., Muschel, R., Humphrey, T., et al. (2022). Ionizing radiation drives key regulators of antigen presentation and a global expansion of the immunopeptidome. Molecular & Cellular Proteomics, 21(11).

Tan, K. M., Petersen, A., and Witten, D. Classification of rna-seq data. In Statistical analysis of next generation sequencing data, pages 219–246. Springer, (2014).

Todd, E. V., Black, M. A., and Gemmell, N. J. (2016). The power and promise of rna-seq in ecology and evolution. Molecular ecology, 25(6):1224–1241.

Toffali, L., D’Ulivo, B., Giagulli, C., Montresor, A., Zenaro, E., Delledonne, M., Rossato, M., Iadarola, B., Sbarbati, A., Bernardi, P., et al. (2023). An isoform of the giant protein titin is a master regulator of human t lymphocyte trafficking. Cell Reports, 42(5).

Upadhyay, G. (2019). Emerging role of lymphocyte antigen-6 family of genes in cancer and immune cells. Frontiers in immunology, 10:819.

Velalopoulou, A., Karagounis, I. V., Cramer, G. M., Kim, M. M., Skoufos, G., Goia, D., Hagan, S., Verginadis, I. I., Shoniyozov, K., Chiango, J., et al. (2021). Flash proton radiotherapy spares normal epithelial and mesenchymal tissues while preserving sarcoma response. Cancer research, 81(18):4808–4821.

Viana, R. S. S., Florentino, H. d. O., Lima, E. A. B. d. F., Fonseca, P. R. d., and Homem, T. P. D. (2011). Heterogeneity correction in the construction of optimized planning in radiotherapy using linear programming. Pesquisa Operacional, 31:565–578.

Wan, T., Wang, H., Gou, M., Si, H., Wang, Z., Yan, H., Liu, T., Chen, S., Fan, R., Qian, N., et al. (2020). Lncrna heih promotes cell proliferation, migration and invasion in cholangiocarcinoma by modulating mir-98-5p/hectd4. Biomedicine & Pharmacotherapy, 125:109916.

Wang, H., Liu, B., and Wei, J. (2021). Beta2-microglobulin (b2m) in cancer immunotherapies: Biological function, resistance and remedy. Cancer letters, 517:96–104.

Wang, L., Rivas, R., Wilson, A., Park, Y. M., Walls, S., Yu, T., and Miller, A. C. (2023). Dose-dependent effects of radiation on mitochondrial morphology and clonogenic cell survival in human microvascular endothelial cells. Cells, 13(1):39.

Wei, T. and Simko, V. R package ‘corrplot’: Visualization of a Correlation Matrix, (2024). URL https://github.com/taiyun/corrplot. (Version 0.95).

Wold, S., Esbensen, K., and Geladi, P. (1987). Principal component analysis. Chemometrics and intelligent laboratory systems, 2(1-3):37–52.

Wu, N., Wang, Y., Wang, K., Zhong, B., Liao, Y., Liang, J., and Jiang, N. (2022). Cathepsin k regulates the tumor growth and metastasis by il-17/ctsk/emt axis and mediates m2 macrophage polarization in castration-resistant prostate cancer. Cell death & disease, 13(9):813.

Xia, K., Huang, X., Zhao, Y., Yang, I., and Guo, W. (2024). Serpinh1 enhances the malignancy of osteosarcoma via pi3k-akt signaling pathway. Translational Oncology, 39:101802.

Xu, P., Ji, X., Li, M., and Lu, W. (2023). Small data machine learning in materials science. npj Computational Materials, 9(1):42.

Xu, S., Hu, E., Cai, Y., Xie, Z., Luo, X., Zhan, L., Tang, W., Wang, Q., Liu, B., Wang, R., et al. (2024). Using clusterprofiler to characterize multiomics data. Nature Protocols, pages 1–29.

Yan, W., Hu, W., Song, Y., Liu, X., Zhou, Z., Li, W., Cao, Z., Pei, W., Zhou, G., and Hu, G. (2024). Differential network analysis reveals the key role of the ecm-receptor pathway in α-particle-induced malignant transformation. Molecular Therapy-Nucleic Acids, 35(3).

Ye, F., Niu, X., Liang, F., Dai, Y., Liang, J., Li, J., Wu, X., Zheng, H., Qi, T., and Sheng, W. (2023). Rnf213 loss-of-function promotes pathological angiogenesis in moyamoya disease via the hippo pathway. Brain, 146(11):4674–4689.

Yin, X., Zeng, D., Liao, Y., Tang, C., and Li, Y. (2024). The function of h2a histone variants and their roles in diseases. Biomolecules, 14(8):993.

Yoo, B. C., Kim, K.-H., Woo, S. M., and Myung, J. K. (2018). Clinical multi-omics strategies for the effective cancer management. Journal of proteomics, 188:97–106.

Yoshioka, N., Kurose, M., Yano, M., Tran, D. M., Okuda, S., Mori-Ochiai, Y., Horie, M., Nagai, T., Nishino, I., Shibata, S., et al. (2022). Isoform-specific mutation in dystonin-b gene causes late-onset protein aggregate myopathy and cardiomyopathy. Elife, 11:e78419.

Yu, J., Deng, X., Lin, X., Xie, L., Guo, S., Lin, X., and Lin, D. (2024). Dst regulates cisplatin resistance in colorectal cancer via pi3k/akt pathway. Journal of Pharmacy and Pharmacology, page rgae104.

Yu, L., Fernandez, S., and Brock, G. (2017). Power analysis for rna-seq differential expression studies. BMC bioinformatics, 18:1–9.

Zannella, V. E., Cojocari, D., Hilgendorf, S., Vellanki, R. N., Chung, S., Wouters, B. G., and Koritzinsky, M. (2011). Ampk regulates metabolism and survival in response to ionizing radiation. Radiotherapy and Oncology, 99(3):293–299.

Zhang, Y., Yuan, Y., Jiang, L., Liu, Y., and Zhang, L. (2023). The emerging role of e3 ubiquitin ligase rnf213 as an antimicrobial host determinant. Frontiers in Cellular and Infection Microbiology, 13:1205355.

Zhao, L., Pridgeon, J. W., Becnel, J. J., Clark, G. G., and Linthicum, K. J. (2009). Mitochondrial gene cytochrome b developmental and environmental expression in aedes aegypti (diptera: Culicidae). Journal of medical entomology, 46(6):1361–1369.

Zhou, G.-Z., Sun, Y.-H., Shi, Y.-Y., Zhang, Q., Zhang, L., Cui, L.-Q., and Sun, G.-C. (2021). Anxa8 regulates proliferation of human non-small lung cancer cells a549 via egfr-akt-mtor signaling pathway. Molecular Biology, 55(5):763–772.

Zhu, L., Jiang, M., Wang, H., Sun, H., Zhu, J., Zhao, W., Fang, Q., Yu, J., Chen, P., Wu, S., et al. (2021). A narrative review of tumor heterogeneity and challenges to tumor drug therapy. Annals of Translational Medicine, 9(16).

